# An integrated pipeline and multi-model graphical user interface for accurate nano-dosimetry

**DOI:** 10.1101/2021.08.31.458389

**Authors:** Ermes Botte, Pietro Vagaggini, Ilaria Zanoni, Davide Gardini, Anna Luisa Costa, Arti Ahluwalia

## Abstract

Accurate dosing of nanoparticles is crucial for risk assessment and for their safe use in medical and other applications. Although it is well-known that nanoparticles sediment, diffuse and aggregate as they move through a fluid, and that therefore the effective dose perceived by cells may not necessarily be that initially administered, dose quantification remains a challenge. This is because to date, methods for accurate dose estimation are difficult to implement, involving precise characterization of the nanomaterial and the exposure system as well as complex mathematical operations. Here we present a pipeline for accurate nano-dosimetry of engineered nanoparticles on cell monolayers, based on an easy-to-use graphical software - DosiGUI - which integrates two well-established particokinetic and particodynamic models. DosiGUI is an open source tool which was developed to facilitate nano-dosimetrics. The pipeline includes methods for determining the stickiness index which describes the affinity between nanoparticles and cells. Our results show that accurate estimations of the effective dose cannot prescind from rigorous characterization of the stickiness index, which depends on both nanoparticle characteristics and cell type.

## Introduction

The unique physicochemical properties of nanomaterials (NMs) are of interest for a wide-range of applications [1], [2]. For this reason, several NMs are now produced industrially and are referred to as engineered nanomaterials (ENMs). For instance, ENMs are extensively employed in the form of engineered nanoparticles (ENPs) for modulating mechanical characteristics of scaffolds, delivering drugs and genes, and labelling in tissue engineering, or as nano-transducers for directing cell behaviour in targeted therapy [3]– [5].

However, the very same nanoscale-related features that make nanoparticles (NPs) - be they natural, incidental or ENPs - so attractive confer potential toxicity to NMs [2], [6]. For this reason, an essential point in nanotoxicology is the characterization of biological effects induced in human tissues and organs as a function of NP dose, along with the investigation of mechanisms triggering their cytotoxicity. Cell cultures *in vitro* represent one of the most common methods for dose assessment, although the extrapolation of dose-response behaviour to the *in vivo* context is still a challenge [7], [8].

A proper definition of the cytotoxic “potential” of a specific NP would require correlating harmful effects to the cumulative amount of material effectively interacting with cells during the exposure test, or what we refer to as the “effective dose” [9], [10]. Cells in culture are known to be sensitive to NPs (and their dissolved ions) that are internalized or even in their immediate vicinity. The cytotoxic effects depend on several intrinsic factors, for example NP size distribution and their effective physical density. Nanotoxicity is also conditioned by extrinsic (*e*.*g*., experimental set-up geometry, cell monolayer uptake) factors which determine the rates of physicochemical phenomena occurring in the system [11]. Although the mechanisms by which NPs induce cellular damage have yet to be fully understood (*e*.*g*., uptake kinetics, oxidative stress) [12], [13], there is plenty of evidence to suggest that only a fraction of the total amount of material initially administered within the suspension (*i*.*e*. the nominal dose) is delivered to cells, representing the effective dose [9], [14]. Moreover, the latter is strongly dependent on the time of exposure and the geometry of the experimental set-up. Due to experimental difficulties in measuring the effective dose, most reports in the literature of *in vitro* dosimetry and nanotoxicology refer to the administered dose, which remains constant in time and is independent of the exposure configuration. Thus, toxicity may be underestimated (*i*.*e*. biological effects are associated with doses higher than those which effectively cause them), making it difficult to properly assess NP hazard and set constraints necessary for the safe use of ENPs [11], [14].

Given this background, *in silico* models are a crucial tool for estimating the effective amount of NPs reaching tissues and cells in specific configurations and performing a more accurate dose-response characterization [9], [10]. In particular, two models represent the state-of-the-art in this field: the *in vitro* sedimentation, diffusion, dissolution and dosimetry (ISD3) model and the distorted grid (DG) model [15], [16].

Prior to using the models, a systematic physicochemical characterization of NPs of interest as well as the identification of unknown extrinsic parameters (*i*.*e*. those determining the uptake kinetics of cells) are essential [17]. In fact, both models require a number of physicochemical characteristics of NPs and the adsorptive or “stickiness” properties of the cell monolayer as inputs. The ISD3 and DG models are distributed as MATLAB codes, requiring full installation of a licensed computing environment and basic programming knowledge for entering input parameters and obtaining dose predictions. As a result, the valuable support that computational models could offer to *in vitro* dosimetry is unexploited, and benefits to NM hazard assessment are limited.

To promote the routine use of NP dosimetry models for accurate prediction of effective dose in *in vitro* systems, we have developed an *in vitro*-*in silico* pipeline leveraged on a graphical user interface (DosiGUI) which integrates the ISD3 and DG models. DosiGUI is an open-source standalone application and allows standardizing the input and output datasets for both models, facilitating their comparison and enabling the identification of the most suitable one for a given NM and experimental configuration. The pipeline includes methods for validating the models through the use of reference highly adsorptive (very sticky) and non-adsorptive (reflective) surfaces for different ENPs. Following this pipeline, we evaluated DosiGUI’s performance for insoluble ENPs by fitting predictions on data generated from experiments performed first with a reflective and maximally adsorptive bottom. Finally, as a proof-of-concept, we applied the pipeline to estimate the effective dose of three different types of ENMs perceived by HepG2 cells in a standard exposure scenario.

## Materials and methods

### The DosiGUI-based pipeline

As depicted in Figure 1, DosiGUI is an essential part of a broader *in vitro*-*in silico* pipeline. The first step of the pipeline is the characterization of NMs. Both the ISD3 and the DG models require NM physicochemical properties as quantitative inputs to perform predictions (Figure 1A). Indeed, NP characteristics such as size, agglomeration and effective density have a major impact on effective dose and must be measured properly. If they are not determined *a priori*, the accuracy of dosimetry calculations may be compromised. Methods for characterizing NPs are extensively reported in the literature, well-established and routinely used [17]–[20]. Extrinsic parameters typical of the specific *in vitro* configuration are also needed (Figure 1B). While some of these features are known by design (*e*.*g*., suspension height, exposure time), those defining the adsorption kinetics (a stickiness index of the bottom boundary) depend on both the properties of the NP and the cell type and should be identified for optimizing the predictive power of the models. Once the input dataset is complete, the models can be reliably run within DosiGUI for simulating the dynamics of NPs and ions in an *in vitro* set-up (Figure 1C). In many cases, simulation results can themselves be used for estimating parameters of the *in vitro* system which may otherwise be difficult to measure, such as the empirical constants describing the affinity between NPs and cells (see the subsection *Identification of stickiness parameters for insoluble ENPs interacting with HepG2 cells*).

**Figure 1.**
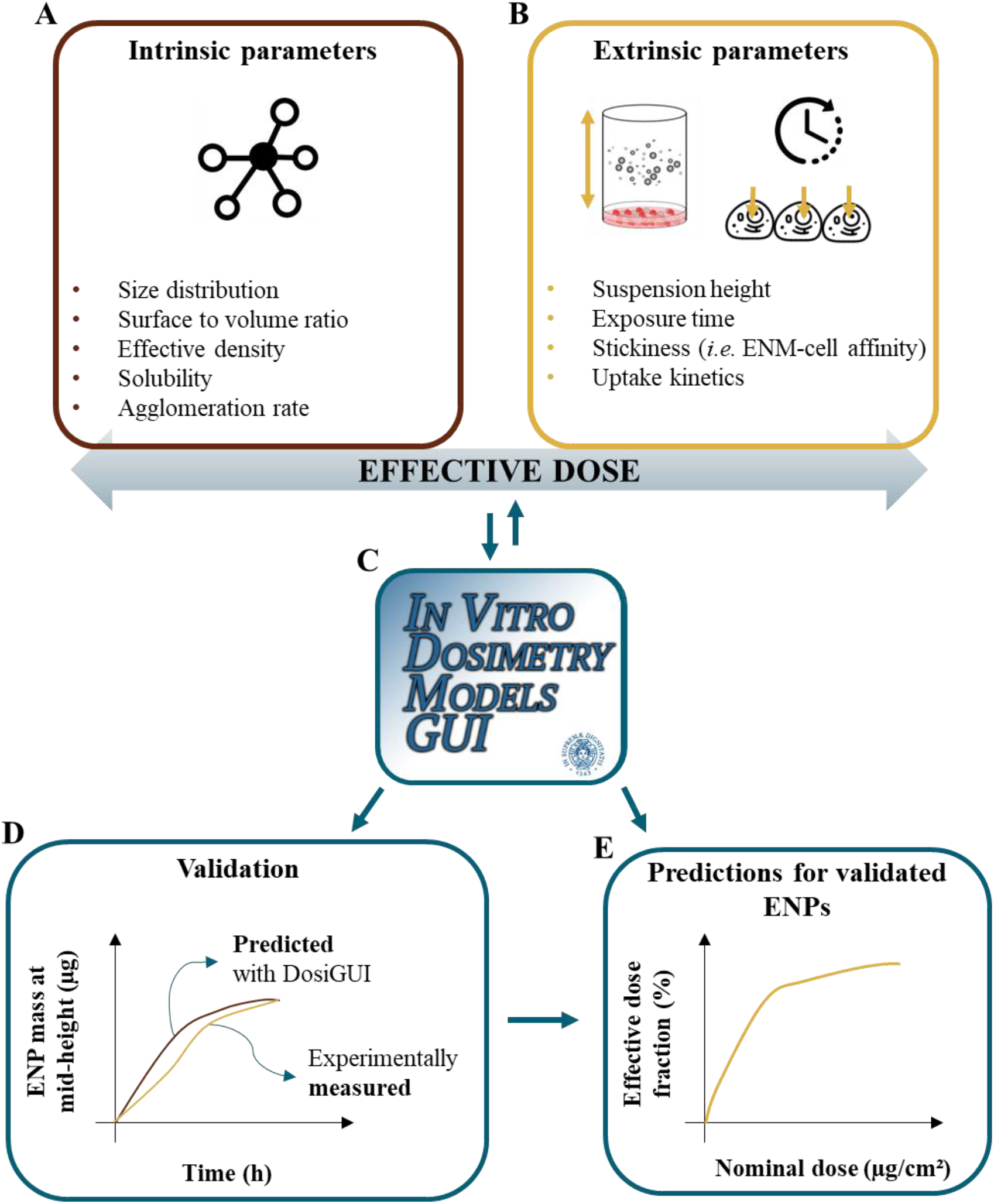
The *in silico*-*in vitro* pipeline leveraging on DosiGUI. Both **A)** physicochemical characterization of ENPs and **B)** extrinsic parameters are crucial for optimizing the reliability of effective dose predictions and improving dose-response characterization. **C)** The complete input dataset is fed to DosiGUI for running simulations, and the outcome can be also exploited for refining extrinsic parameters (*e*.*g*., the cell layer stickiness). **D)** DosiGUI is validated for specific ENPs by comparing predictions and experimental measurements of a fully characterized test system. **E)** Once validated, DosiGUI predictions of effective dose for a given ENP can be extrapolated to various exposure scenarios.

Given a specific NP, the validation of DosiGUI predictions (Figure 1D) consists in comparing the experimentally measured deposited mass over time in a fully characterized test system with the corresponding profile estimated by simulating the chosen set-up *in silico* (see the subsection *DosiGUI validation for three insoluble ENPs*). If there is a statistically significant correlation between the two, the model(s) can be assumed valid and hence used to predict the effective dose of the given NP in different exposure scenarios (Figure 1E).

### DosiGUI: a multi-model graphical user interface for in vitro nano-dosimetry

DosiGUI was developed as an open source, standalone desktop application. We implemented it in the MATLAB computing environment (R2020b, The MathWorks^®^ Inc., Boston, Massachusetts) by means of the App Designer Toolbox. Its source code was compiled to run as a desktop application for both Windows and iOS operating systems, without MATLAB. When DosiGUI is installed, MATLAB Runtime (version 9.9 for DosiGUI) is automatically downloaded and run. A practical guide on how to download and install DosiGUI is provided in the SI (section SI1).

Once the installation is complete, launching the application from the desktop shortcut opens a main window where a brief description of ISD3 and DG is provided (Figure 2A). After initiating one of the two models by clicking on the corresponding *Start* button, the user can manually enter all input parameters needed for running simulations. Alternatively, a pop-up menu (Figure 2B) allows the input parameters to be either directly loaded from a specific file containing the information requested for the chosen model (*Load from file*) or obtained by automatic rearrangement of an input file which may have been already stored for the other model (in Figure 2B, *Load DG simulation*). A single harmonized input dataset which is suitable for both models is one of the unique features of DosiGUI: the same experimental configuration can be simulated using ISD3 and DG, and the consistency of their predictions can be compared to determine the most suitable model for estimating the effective dose of NPs for a given set-up. As an example, Figure 2B show the interface panels for ISD3 and all categories of inputs required by the model, including intrinsic and extrinsic parameters (left side). Specific plot settings are also listed on the right: predicted effective doses as well as any other available output variable can be plotted as a function of time or height, and the generated figures can be saved as .png, .jpg, .tif or .pdf files. Once started, a status bar shows the current status of the simulation. When the simulation is complete, inputs and outputs are saved as .xlsx and .mat files. Finally, all calculations performed by DosiGUI are summarized and saved into a .txt file named *logfile*. The logfile helps in identifying interruptions caused by computational errors when the GUI is running.

**Figure 2.**
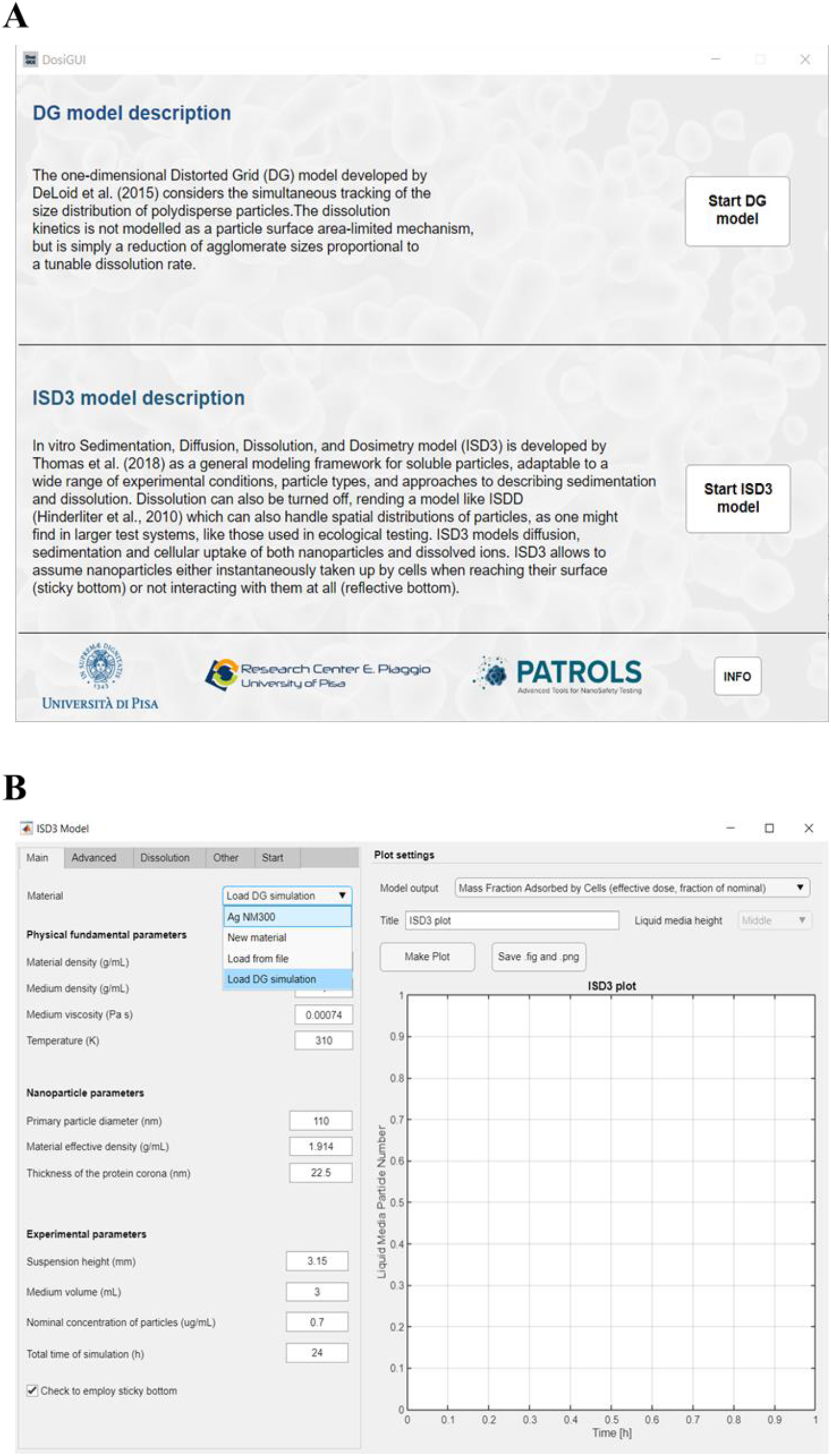
The graphical user interface. **A)** The main window of DosiGUI. **B)** The interface panel for the ISD3 model. The input values shown refer to the validation of ISD3 for silver (Ag - NM-300K) nano-dosimetry, carried out in [10].

#### Basic principles of integrated dosimetry models

The ISD3 and DG models simulate the temporal and spatial dynamics of NPs suspended in cell culture medium, in a container with constant cross-sectional area. They both consider a one-dimensional (1D) model of sedimentation, diffusion and, if applicable, dissolution with an adsorptive bottom surface, which mimics the presence of a cell monolayer [15], [16]. The different processes are described through a series of rate equations (reported in SI2), which are solved numerically to give NP diameter (*d*_*p*_ (m)) and number surface density (*N* (m^-2^)) as functions of time (*t* (s)) and the 1D space dimension (*x* (m), corresponding to height). They differ in: i) the method of solving equations, ii) the way they treat dissolution and iii) the boundary layer condition, which basically describes the adsorptive properties of the bottom. In particular, the DG model accounts for NP dissolution considering a first order rate equation with a rate constant that depends on *d*_*p*_. On the other hand, the ISD3 implements dissolution as a surface area-driven phenomenon based on a NP-specific kinetic model. This description can be more accurate but less general than that provided by the DG model. It does however require an in-depth characterization of dissolution kinetics for determining an analytical formulation of the associated rate suitable for the NP of interest.

As far as the bottom boundary condition (*i*.*e*. the cell monolayer adsorption) is concerned, both models provide a tuneable stickiness to be set through a parameter, which describes adhesive behaviour ranging from a purely reflective condition up to a total instantaneous adsorption. The DG model accounts for this by means of a Langmuir isotherm adsorption, expressed as in Eq. (1):

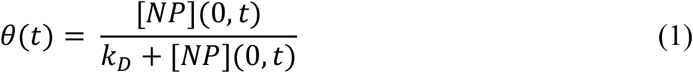

where *θ* is the surface fraction of the bottom boundary occupied by adsorbed NPs and [*NP*](0, *t*) (mol L^-1^) is the molar concentration of NPs in the vicinity of the boundary. *k*_*D*_(mol L^-1^) - the parameter to be set for modulating the bottom boundary stickiness - denotes the equilibrium dissociation constant; it is inversely proportional to the NP-cell affinity or stickiness index. The ISD3 deals differently with the bottom adsorption, implementing Eq. (2).

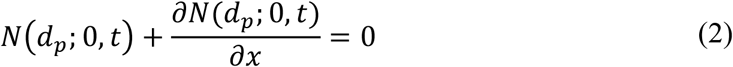

This is the typical formulation of a boundary condition for the partial differential equation (PDE) solved by the model. As detailed in SI3, we have modified Eq. (2) to encompass the entire range of possible stickiness conditions, from the totally reflective (no adsorption) to the maximally adsorptive (instantaneous uptake) boundary, through a parameter *K* (m) which is inversely proportional to the stickiness index. Methods for the identification of parameters describing the stickiness index of the bottom boundary and their application to a specific exposure set-up are reported in the subsection *Identification of stickiness parameters for insoluble ENPs interacting with HepG2 cells*.

Intrinsic and extrinsic parameters constitute the input dataset requested by DosiGUI to run simulations. In Table 1, we provide a list of the most important parameters which impact the outcome of the models.

**Table 1.** Basic intrinsic and extrinsic input parameters required by DosiGUI for running the ISD3 and DG models.

| ENM intrinsic parameters | Extrinsic (configuration dependent) parameters |
| --- | --- |
| ENP size distribution (*i.e.* existing diameters ($d_p$) and corresponding normalized counts) | Medium height ($h$) |
| Density ($\rho$) | ENM nominal dose ($M_{nom}$) |
| Effective density ($\rho_{eff}$) | Bottom boundary stickiness ($k_D$ or $K$) |
| Dissolution constants<sup>*</sup> | Total exposure time ($\tau$) |
| Medium density ($\rho_f$) | Spatial resolution ($\Delta x$) |
| Medium dynamic viscosity ($\mu$) | Time resolution ($\Delta t$) |
| Medium temperature ($T$) | |

<sup>\*</sup>Depending on the kinetic model adopted for dissolution in the chosen computational framework

In this work, for the proof-of-concept application of the approach, we focused on demonstrating the rigorousness of our nano-dosimetry pipeline for three referenced insoluble ENPs: titanium oxide (TiO_2_ - NM-105), cerium oxide (CeO_2_ - NM-212) and barium sulphate (BaSO_4_ - NM-220). The input physicochemical datasets were taken from Keller *et al*. [19], since the same ENPs and dispersion protocols were employed in this study.

### DosiGUI validation for three insoluble ENPs

The first step towards accurate nano-dosimetry and the construction of effective dose-response curves consists in model validation. To this end, we set up a test system to measure ENP sedimentation simulating the two extreme boundary conditions (*i*.*e*. reflective bottom and maximal adsorption). The experimental configurations were implemented in both ISD3 and DG within DosiGUI for providing computational predictions and evaluating their fitting to the corresponding *in vitro* data. All statistical analyses were carried out in GraphPad Prism (v7, GraphPad Software^®^).

#### Experimental measurements of mid-height mass fraction

All the evaluated ENPs were purchased from JRC. Standard cell culture medium (DMEM), bovine serum albumin (BSA) and fetal bovine serum (FBS) - required for preparing the suspension - were supplied by ThermoFisher. For the preparation of the adsorptive gel, gelatin (Sigma-Aldrich) cross-linked with GPTMS (AlfaAesar) was used.

ENP stocks were prepared dispersing the nano-powders in MilliQ water and 0.05% w/v of BSA, following the Nanogenotox protocol [21]. Samples were prepared diluting the stock in DMEM with 10% w/v FBS at mass concentrations of 5, 25 and 50 mg L^-1^ (corresponding to nominal doses of 1.5, 7.8 and 15.6 µg cm^-2^, respectively). 8 mL of dispersion were put in plastic vials and incubated for 1, 4 and 24 h (5% CO_2_, 37 °C, 95% RH). The 8 mL were chosen because they reproduce the geometry of a 96-well plate on a bigger scale. To mimic the condition of maximal adsorption, we coated the bottom of the vials with a thick layer of sticky gel (gelatin 1% w/v cross-linked with 100 µL of GPTMS per gram of gelatin, see SI for further details); for the reflective boundary condition, the vial bottom was left untreated. In both cases, after the incubation time, a volume of 3 mL (*V*_*sam*_) was collected from the middle of the suspension column and analysed. The sampling procedure was manual, according to a specific protocol validated by ISTEC laboratories (see SI). All experiments were carried out in triplicate.

The elemental composition of the dispersion was assessed by inductively coupled plasma optical emission spectrometry (ICP-OES), using an ICP-OES 5100 – vertical dual view apparatus coupled with OneNeb nebulizer (Agilent Technologies, Santa Clara, CA, USA). Before measuring the associated ENP mass concentration, samples were acidified with 0.3 mL of nitric acid (HNO3 65%, Sigma-Aldrich - St. Louis, MI, USA). Calibration curves were obtained with standard samples containing 0.05, 0.10, 1.00, 10.00 and 100.00 mg L^-1^ of each ENP suspended in DMEM with FBS 10% w/v and after applying the same digestive procedure.

Then, for each analysed sample, Eq. (3) was used to convert the outcome of the measurement from a mass concentration to a fraction of the initially administered ENM mass 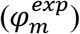:

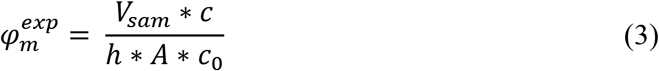

where *c*_0_(µg mL^-1^) is the nominally administered mass concentration, *c* (µg mL^-1^) denotes the mass concentration detected by ICP-OES within the sampled volume *V*_*sam*_at mid-height, *h* (mm) is the total height of the liquid column and *A* (mm^2^) is the cross-sectional area of the vial. Given that the mid-height and the initial concentrations refer to different volumes, they have to be rescaled as absolute masses to estimate the ENP fraction contained in *V*_*sam*_. Equivalently, since *A* is constant along the liquid column, the ENP fraction can be expressed as the ratio of corresponding values of dose (*i*.*e*. masses per unit area as a function of time and height, see SI2).

#### In silico predictions of mid-height mass fraction and correlation analysis

To model the *in vitro* experiments, the dynamics of ENPs along the liquid column was simulated separately running the models embedded within DosiGUI on a workstation equipped with an 8^th^ generation Core™ i7 (Intel Corporation^®^) microprocessor. The maximally sticky boundary condition in the presence of gelatin was taken into account considering *K* = 0 m for ISD3 and *k*_*D*_ = 0 mol L^-1^ for DG, while a purely reflective bottom was replicated by setting relatively high values for both parameters (*K* = 10^12^ m and *k*_*D*_ = 10^−5^ mol L^-1^), such that further increasing them does not imply any further reduction of the adsorption. Dissolution was disabled, given the negligible solubility in aqueous media of the three ENPs. The spatial and temporal resolutions were set respectively to 0.5 mm and 1 min to optimize the accuracy of predictions.

The results of simulations in terms of ENP mass along the liquid column at the time points of interest (*i*.*e*. 1, 4 and 24 h) were rearranged according to Eq. (4), to obtain the fraction of the initially administered mass 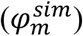 contained in *V*_*sam*_.

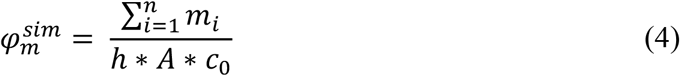

In the equation, *m*_*i*_ (µg) is the ENP mass in the i-th simulation element at the considered time point, and *n* = *V*_*sam*_/∆*x* * *A* denotes the number of simulation elements in *V*_*sam*_.

Then, we evaluated the reliability of simulated data from both ISD3 and DG through correlation analysis between measured and predicted mid-height mass fraction profiles over time (see SI4 for details on the statistical analyses).

### Identification of stickiness parameters for insoluble ENPs interacting with HepG2 cells

After the goodness of model predictions for NM-105, NM-212 and NM-220 was assessed, we used DosiGUI to estimate the effective dose delivered to HepG2 cells in a standard exposure configuration. As discussed previously, a crucial parameter for the model is the stickiness index. In fact, when adsorbed by cells, the ENPs are effectively removed from the suspension; consequently, the downward diffusive flux is increased and this may significantly impact the effective dose [16]. The stickiness index for each ENP was evaluated through estimation of the empirical constants *K* and *k*_*D*_ (see Eq.s (1) and (2)). The experiments consisted in measurements of the quantity of ENP mass adsorbed by HepG2 over time. The data were compared through correlation analysis with their corresponding computational predictions, obtained through parametrization of *K* or *k*_*D*_. Using this method, we were able to identify the most suitable HepG2-ENP stickiness index for the three ENPs used.

#### Experimental measurements of cell adsorbed mass fraction

HepG2 cells were seeded to confluence in 96-well plates (about 10^5^ cells/well) and incubated overnight (5% CO_2_, 37 °C, 95% RH). ENPs were suspended in DMEM, with 10% w/v BSA. Suspensions containing 25, 50 and 250 µg mL^-1^ of each ENP (corresponding to nominal doses of 7.8, 15.6 and 78.1 µg cm^-2^, respectively) were obtained through serial dilutions of a stock (2.56 mg mL^-1^). Then, a working volume of 100 µL of suspension was added to wells in four different plates, one for each exposure time (*i*.*e*. 4, 8, 24 and 72 h). After incubation, the ENP suspension was removed, and wells rinsed twice with PBS 1X for eliminating ENPs either deposited (but not adsorbed) onto cells or attached to the lateral wall of the well. Trypsin-EDTA was then used for detaching cells, and samples were collected for measuring the adsorbed ENP mass. The overlaying ENP suspension together with the washing volume of PBS was also stored and tested to account for all the initially administered mass. Blank controls (BCs) consisted in cells without ENPs, while positive controls (PCs) were simply 100 µL of the suspension, collected immediately after the stock dilution. All experiments were carried out in triplicate.

All the collected samples were diluted in MilliQ water to a total volume equal to *V*_*sam*_. Then, the same protocol described for validation was applied for performing ICP-OES measurements, determining the ENP mass concentration in each sample. Note that, for each ENP and nominal dose, BCs and PCs represent the offset of the measurement and the maximum mass concentration detectable in the samples, respectively. Thus, to get meaningful adsorbed fractions 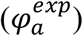 the raw data were post-processed as follows:

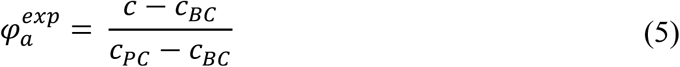

where *c* (µg mL^-1^) is the adsorbed ENP concentration measured by ICP-OES, while *c*_*BC*_ and *c*_*PC*_(µg mL^-1^) are the ENP concentrations detected for the corresponding BCs and PCs, respectively. All the terms in Eq. (5) are expressed as mean ± standard deviation of the triplicate, and standard methods were applied to account for error propagation. Since they all refer to the same volume *V*_*sam*_, the adsorbed fraction 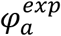 refers to mass as well as concentration.

#### In silico predictions of cell adsorbed mass fraction and correlation analysis

As reported for validation, exposure tests for each ENP were replicated *in silico* by running both models in DosiGUI. After setting the input parameters (geometry and ENP characteristics), a parametric sweep was implemented for the stickiness index by iteratively varying *K* and *k*_*D*_ within reasonable ranges (reported in the SI, Table S2). Simulations were carried out for the same nominal doses experimentally administered, setting the spatial and temporal resolution respectively to 0.05 mm and 1 s to optimize the accuracy of predictions.

In this case, to get predictions of adsorbed mass fractions 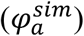 comparable to their experimental counterparts, for each time point of interest the model outcome was rearranged as in Eq. (6):

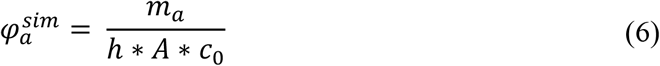

where *m*_*a*_ (µg) is the cumulative ENP mass adsorbed at the bottom of the column, and the other parameters are the same as in Eq. (4).

Once the experimental and simulated datasets were homogenized, we performed a correlation analysis for identifying the most reliable value of both *K* and *k*_*D*_ for each of the three ENPs, *i*.*e*. the HepG2 stickiness with respect to those kinds of ENPs. Specifically, the best fitting regression line for 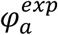 as a function of 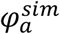 was estimated with a confidence interval of 95% for each model, ENP and nominal dose with respect to each stickiness index considered. The most suitable value of *K* or *k*_*D*_ was determined optimizing the same criteria for the goodness of fit as in the section *DosiGUI validation for three insoluble ENPs*. Details on the statistical analyses are provided in SI4.

### Prediction of the effective dose of insoluble ENPs delivered to HepG2 cells

Once the specific ENP-cell type combination was fully characterized, we were able to reliably extend DosiGUI predictions to any exposure scenario involving that combination. Thus, to complete the proof-of-concept application of the introduced nano-dosimetry pipeline, we chose to estimate the effective dose of each of the three ENPs perceived over time by a HepG2 monolayer using the experimental configuration described for identifying the stickiness index. The optimum value of *K* or *k*_*D*_ identified in the second set of experiments was set for each ENP; all other input parameters were as described in *Identification of stickiness parameters for insoluble ENPs interacting with HepG2 cells*.

## Results

### DosiGUI validation for three insoluble ENPs

Table 2 shows how predicted profiles of mid-height mass fraction over time correlate with experimental data. In particular, values of the correlation statistics considered (see SI4) are shown for the “worst case” of the more predictive model, *i*.*e*. for the nominally administered dose giving the weakest correlation when using the model which results as the most suitable for the specific ENP. Previous reports have demonstrated the robustness of ISD3 for different kinds of NMs (Ag - NM-300K) [10]. However, for the three ENPs studied here, the DG model reliably estimates mass fractions over time at mid-height, since even the worst-case correlation is statistically significant.

**Table 2.** Results of the correlation analysis between measured and simulated mid-height mass fraction profiles over time. The Pearson’s coefficient (*r*) and the p-value of the extra sum-of-squares F-test on the slope of the regression line (*p*) are reported for the “worst case” of the more predictive model (specified in the last column).

| ENP | $r$ | $p$ | Most suitable model |
| --- | --- | --- | --- |
| TiO<sub>2</sub> (NM-105) | 0.987 | 0.534 | DG |
| CeO<sub>2</sub> (NM-212) | 0.826 | 0.696 | DG |
| BaSO<sub>4</sub> (NM-220) | 0.999 | 0.056 | DG |

### Identification of stickiness parameters for insoluble ENPs interacting with HepG2 cells

Table 3 shows optimal values of the parameters determining the stickiness index of HepG2 cells with respect to each of the three ENPs analysed. For both models, the stickiness values obtained differ significantly for each ENP, underlining that the same cell type differentially uptakes NPs based on their physicochemical traits. In Figure 3, the mass fraction adsorbed over time by the HepG2 monolayer for each ENP estimated by the DG model is shown for two cases: i) *k*_*D*_ = 0 mol L^-1^ (maximal stickiness, DG model); ii) *k*_*D*_ values estimated from the ICP-OES data. A similar figure is reported in the SI for ISD3 (Figure S1). The figure highlights the importance of characterizing the boundary stickiness to obtain meaningful estimates of effective dose.

**Table 3.** Optimal values for parameters determining the HepG2 stickiness indices with respect to insoluble ENPs, identified for both the ISD3 (*K*_*opt*_) and the DG model (*k*_*D,opt*_).

| **ENP** | **$K_{opt}$ (m)** | **$k_{D,opt}$ (mol L<sup>-1</sup>)** |
| --- | --- | --- |
| TiO<sub>2</sub> (NM-105) | $8.0 \times 10^6$ | $3.0 \times 10^{-9}$ |
| CeO<sub>2</sub> (NM-212) | $8.0 \times 10^4$ | $2.0 \times 10^{-9}$ |
| BaSO<sub>4</sub> (NM-220) | $6.4 \times 10^4$ | $5.8 \times 10^{-10}$ |

**Figure 3.**
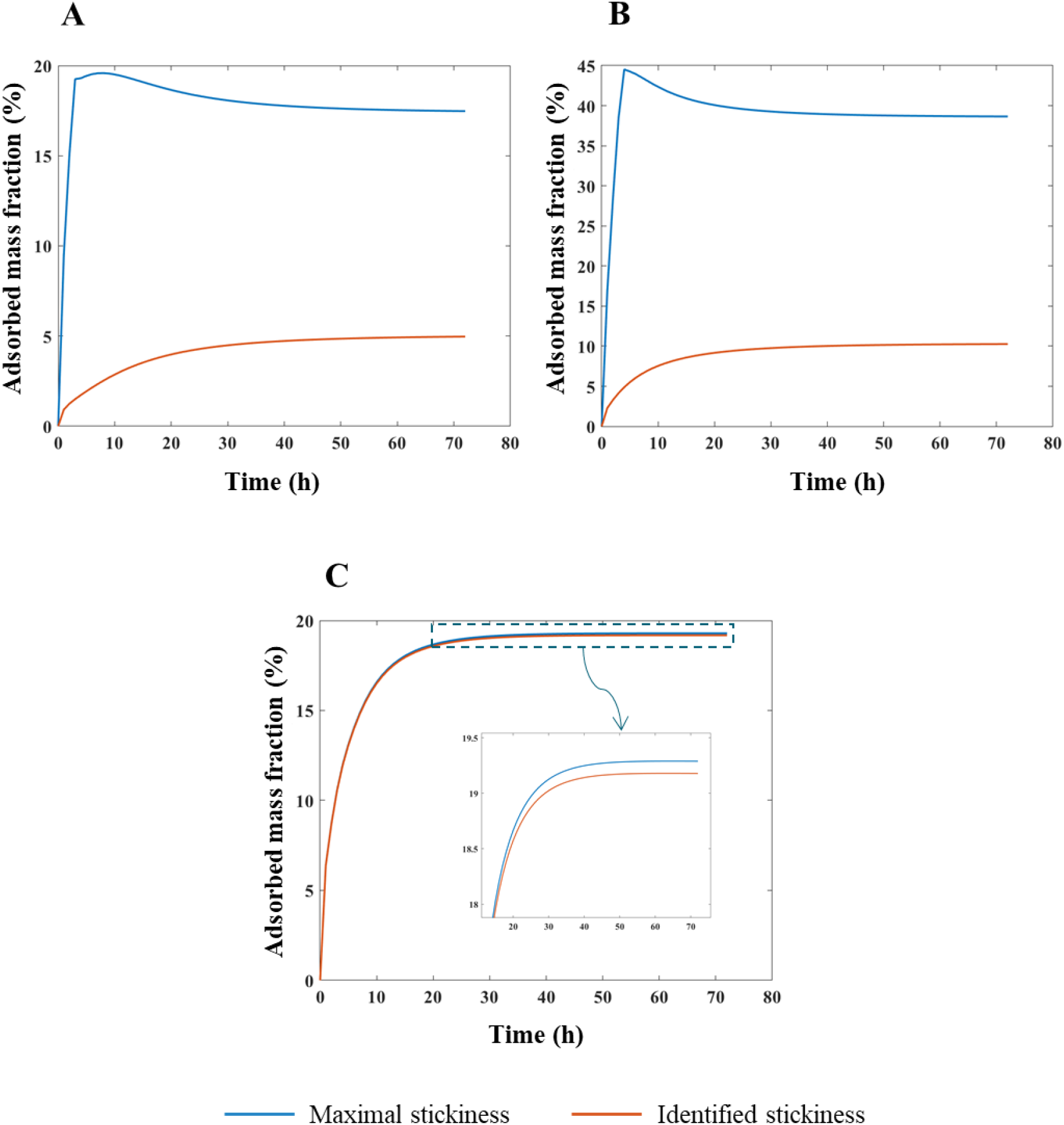
DosiGUI predictions of mass fraction over time adsorbed by HepG2 cells expressed as a percentage of the total amount of ENP mass in the suspension, assuming a nominally administered dose of 78.1 µg/cm^2^. Comparison between the case of a maximally sticky bottom (blue line) and the identified stickiness index of HepG2 monolayers (orange line) for: **A)** TiO_2_ (NM-105), **B)** CeO_2_ (NM-212) and **C)** BaSO_4_ (NM-220).

Using the correct stickiness index results in significantly lower adsorbed mass fractions for NM-105 and NM-212 (Figure 3A and 3B). On the other hand, the two curves in Figure 3C for NM-220 are similar. In fact, as reported in Table 3, this ENP is very highly adsorbed on the HepG2 monolayer, approaching nearly complete stickiness.

### Prediction of the effective dose of insoluble ENPs delivered to HepG2 cells

Figure 4 reports simulated effective doses delivered to HepG2 cells in monolayer, starting from different nominal doses. These results close the loop of the proof-of-concept application of the integrated pipeline proposed here, leveraging on the successful validation of DosiGUI predictions for the insoluble ENPs analysed and exploiting the accurately characterized stickiness indices between ENPs and HepG2 cells. All graphs in the figure refer to simulations carried out with the DG model, since it proved to be the most suitable for replicating the dynamics of the three ENPs. Corresponding predictions by the ISD3 model are reported in the SI (Figure S2).

**Figure 4.**
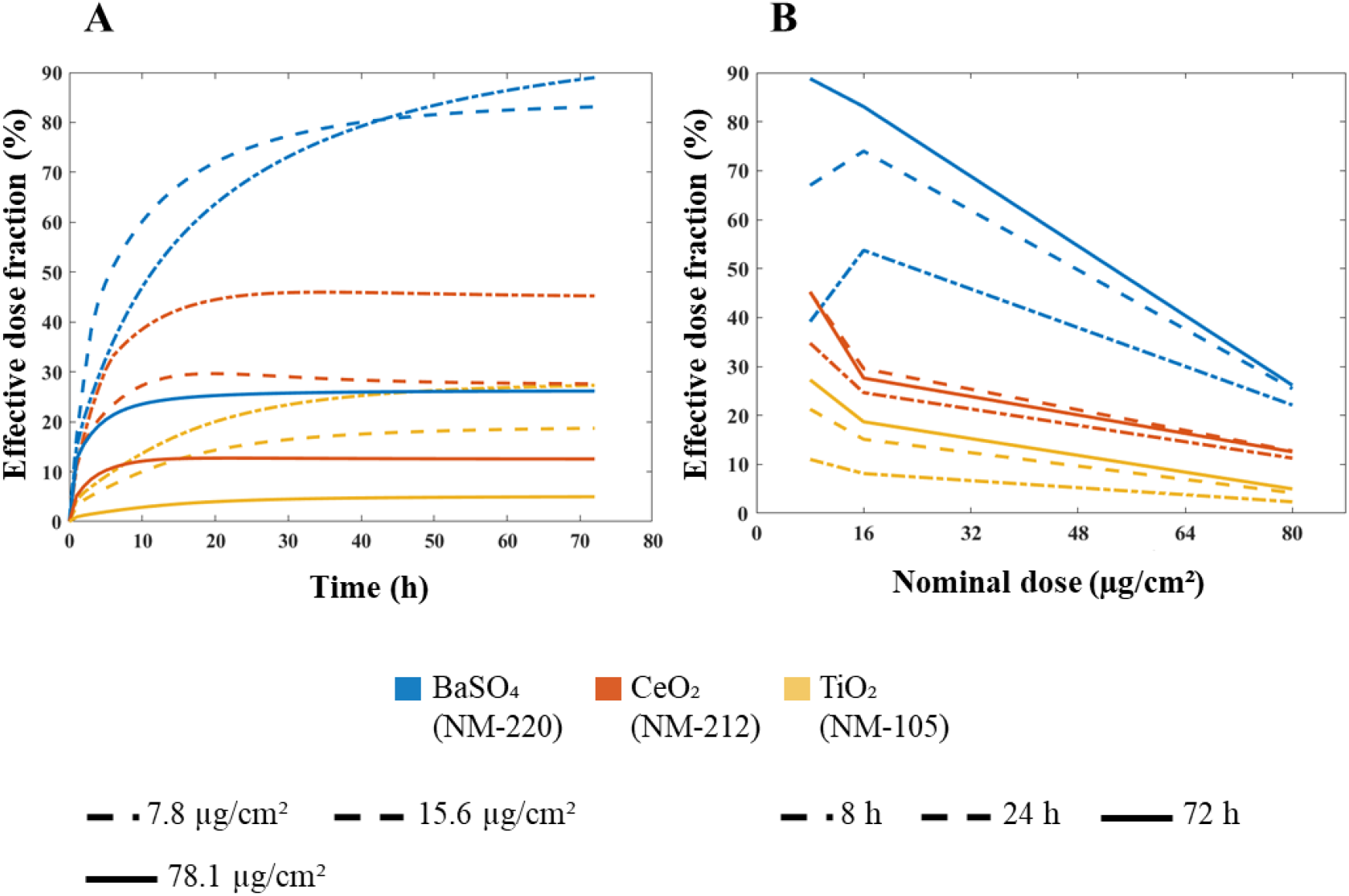
DosiGUI predictions for the effective dose fraction (*i*.*e*. effective dose/nominal dose, percentage) delivered to HepG2 cells. **A)** Effective dose fraction versus exposure time. **B)** Effective dose fraction versus nominal dose.

As expected, cells do not interact with all of ENM administered, even for long exposure times (Figure 4A). On the contrary, a “saturation” effect is observed along with increasing nominal doses, in terms of a reduction of effective dose fraction (*i*.*e*. effective dose/nominal dose) (Figure 4B).

## Discussion

Accurately determining the amount of ENPs that cells, tissues or organs are exposed to in specific scenarios (*i*.*e*. the effective dose) is a key step towards safely tapping the full potential of ENMs in biomedicine. However, due to technical limitations and the complexity of nano-dosimetry, toxicologists still struggle to provide rigorous estimations. This may result in a misleading characterization of NP toxicity and consequent hazard assessment.

To tackle these issues, we present an *in silico*-*in vitro* pipeline for accurately assessing effective doses in monolayer cell cultures. The approach relies on DosiGUI, a purposely developed multi-model graphical user interface. DosiGUI is open source and embeds the two state-of-the-art *in silico* models for computational nano-dosimetry. The pipeline starts with a rigorous characterization of the exposure configuration of interest for the generation of an input parameter dataset. Among the intrinsic characteristics of NPs and the extrinsic parameters of the exposure set-up necessary for running the models, those determining the adsorption kinetics of NPs on cells strongly impact on the effective dose (as highlighted in Figure 3) but have often been neglected. Thus, in this work we introduced new methods for the identification of parameters determining the cell-NP stickiness index.

As a proof-of-concept, we demonstrated the robustness of the approach for three ENPs (NM-105, NM-212, NM-220), which are reported to have a negligible solubility. Starting from physicochemical characterization reported by Wohlleben’s group [19], the reliability of simulations performed using DosiGUI was validated experimentally reproducing and predicting the effective dose for two reference boundaries with a totally adsorptive and a purely reflective surface, respectively. This allows identifying the most suitable model to simulate the dynamics of each ENP. For the three ENPs studied in this work, the DG model was the most suitable (Table 2).

Following this first validation step, parameters describing the adsorption kinetics of ENPs on HepG2 cells were identified so as to set a realistic stickiness index for simulations. The results confirmed that the boundary stickiness is an ENP-specific feature. In fact, the values of kinetic constants for the three ENPs differ significantly from each other, with NM-220 showing the highest affinity for HepG2 (Table 3). Thus, setting the right stickiness index is a crucial step for accurate effective dose estimation. However, in many cases the value of the stickiness index may not be readily available, since the parameter identification procedure is rather complicated and time-consuming. For this reason, we have added a slider which allows setting the stickiness index at five levels (reflective, low, medium, high, maximal), which correspond to a reasonable discretization of the ranges of *K* or *k*_*D*_ considered for the optimization process.

Having characterized the adsorptive behaviour of HepG2 cells, we employed DosiGUI to predict the effective dose of each ENP in a standard *in vitro* test configuration. As expected, only a fraction of the administered amount of ENPs is computed to interact with the monolayer, with a “saturation” effect emerging at high nominal doses (Figures 4 and S2). This is because the adsorption of sedimenting NPs is a surface occupancy-driven mechanism. The mechanism is explicitly modelled in DG (see Eq. (1) and section SI3), which may be the reason why its predictions better correlate with experimental data for insoluble ENPs. As expected, effective doses estimations are higher for ENPs characterized by higher stickiness indices.

In conclusion, this study provides an important contribution to more accurate dose-response characterization of NPs and to the improvement of safety and risk assessment resulting from the exposure of human tissues and organs to commonly employed ENPs. DosiGUI should facilitate the incorporation of *in silico* tools in nanotoxicology, encouraging more data sharing, cross-laboratory comparisons and an exhaustive characterization of cell stickiness for a wide spectrum of phenotypes, which could also be integrated as an essential part of existing ENP databases. Finally, the *in silico* models embedded within DosiGUI could be extended to three-dimensional (3D) configurations for effective dose estimations in *in vitro* cell aggregates having higher levels of complexity, such as spheroids and organoids. Besides improvements in nano-dosimetry, 3D models could be used to simulate the dynamics of NPs in more physiologically relevant scenarios and for a better *in vitro*-to-*in vivo* extrapolation.

## Acknowledgements

This work has received funding from the European Union’s Horizon 2020 Research and Innovation Program under grant agreement No. 7601813 PATROLS project.

