## Supplementary Information for "An integrated pipeline and multi-model graphical user interface for accurate nano-dosimetry"

### SI1 – A practical guide for downloading and installing DosiGUI

The installer of the standalone version of DosiGUI is freely available at this link. Before beginning the installation procedure, ensure you have a stable internet connection; then, follow the steps illustrated below.

1) Download the files attached at the link, by clicking on the green button *Code* and choosing "Download ZIP".

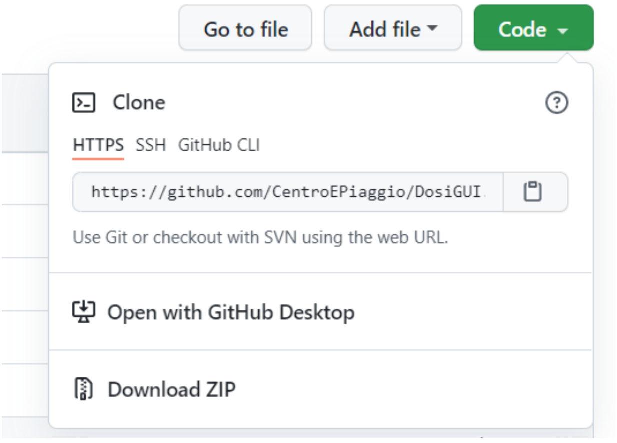


2) Unpack the executable file (*DosiGUI_Web_Installer*) suitable for your operating system. It is the installer of all required components for running DosiGUI as a desktop application. For Mac-users, please refer to the file *MacOS Readme.pdf* for further details, available at the same link in the folder "MacOS".
3) Launch the *DosiGUI_Web_Installer* executable file and wait for the loading window to appear.

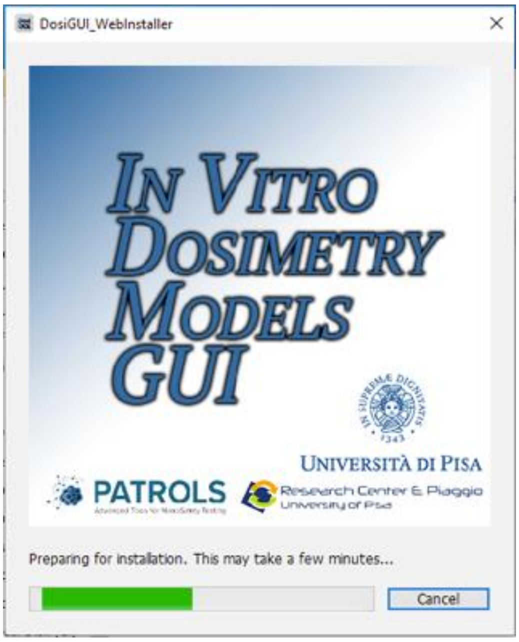


4) Once the loading is complete, a new window appears, where you can check the version of the application you are about to install. Click *Next* to proceed.

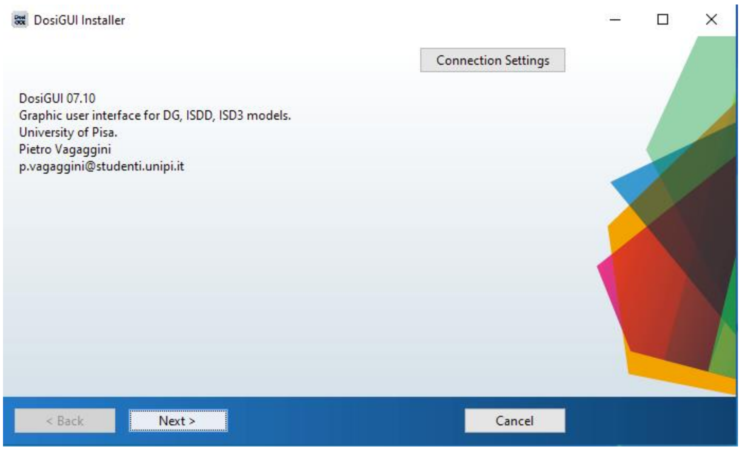


5) Choose the desired folder to install the application. It is recommended to keep the default settings and check the option “Add a shortcut to the desktop”.

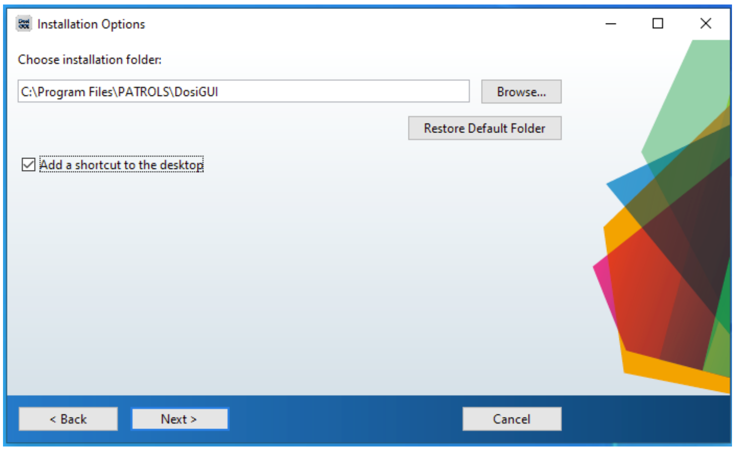


6) Choose the desired folder to install MATLAB Runtime (a free MATLAB component required to run DosiGUI). It is recommended to keep the default settings.

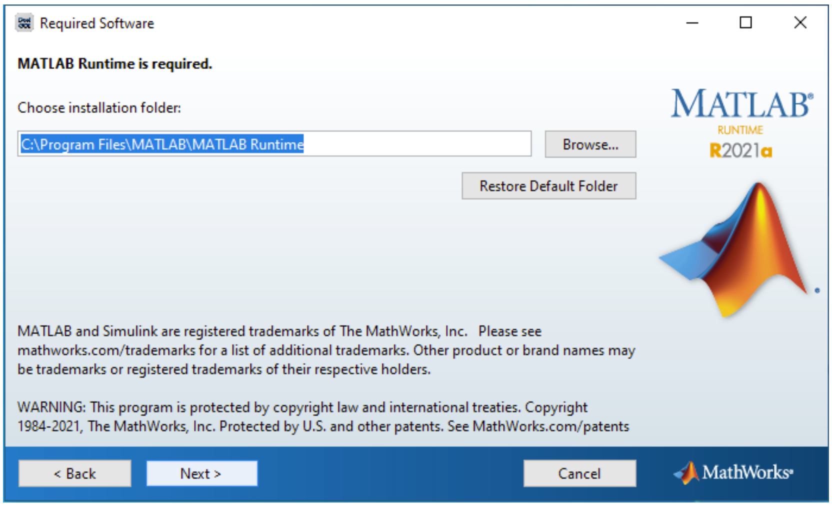


7) Accept the license agreement for MATLAB Runtime and click *Next*.

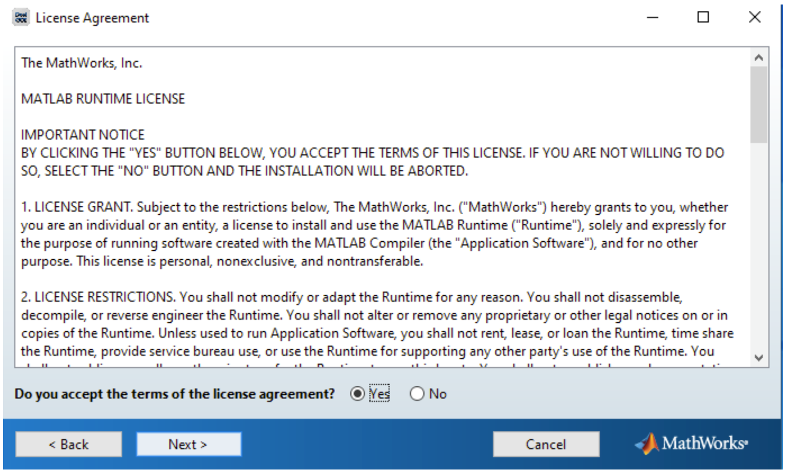


8) A confirmation window summarizing the installation settings appears. Click *Install* to start the installation.

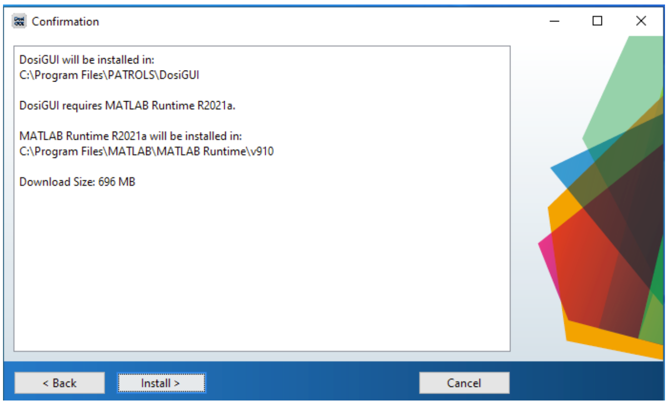


9) The operation begins with the download and installation of MATLAB Runtime (approximately 500MB). Then, DosiGUI is installed.

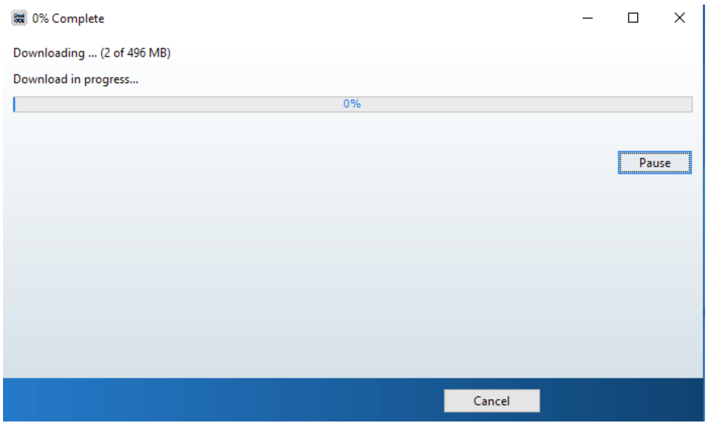


10) Once the installation is complete, close the installer window. Now, you should be able to run DosiGUI from the shortcut on your desktop.

### SI2 – Analytical details about dosimetry models integrated within DosiGUI

Both the ISD3 and DG models refer to the dynamics of NPs suspended in a cylinder containing protein-enriched culture medium which a cell monolayer on the bottom is exposed to [1], [2]. Considering a homogeneously sticky bottom boundary (*i.e.* a homogeneously distributed cell monolayer), requirements for Cartesian symmetry are verified, thus the system can be studied by means of a 1D model representing the axial direction ($x$) only. Along this axis, three physical phenomena are taken into account: sedimentation, diffusion and, if applicable, dissolution, described by the following rate equations.

$$R_{diff}(d_p;x,t) \;=\; D_{diff}(d_p)\frac{\partial^2 N(d_p;x,t)}{\partial x^2} \tag{S1}$$

$$R_{sed}(d_p;x,t) \;=\; -\,v_{sed}(d_p)\frac{\partial N(d_p;x,t)}{\partial x} \tag{S2}$$

$$R_{diss}(d_p;x,t) \;=\; -\,\frac{\partial}{\partial d_p}\left(N(d_p;x,t)\frac{\partial d_p}{\partial t}\right) \tag{S3}$$

The variables and parameters introduced in Eqs. (S1) – (S3) are listed below:

- $x$ (m) is the spatial coordinate;
- $t$ (s) is the temporal coordinate;
- $d_p$ (m) denotes the NP effective diameter (*i.e.* the diameter of the NP and its solution-dependent corona);
- $R_{diff}, R_{sed}$ and $R_{diss}$ ($m^{-2}$ $s^{-1}$) are the diffusion, sedimentation and dissolution rates, respectively;
- $N$ ($m^{-2}$) denotes the surface density of NPs having a certain diameter $d_p$. $N$ is the variable that the models solve for in each spatial point at each iteration and is defined for each NP size in the dispersion. Thus, the corresponding mass surface density field $M$ (kg $m^{-2}$) is:

$$M\big(d_p;x,t\big) = \rho_{eff}\frac{\pi}{6}{d_p}^3 N(d_p;x,t) \tag{S4}$$

with $\rho_{eff}$ (kg m$^{-3}$) the effective physical density of the NP (*i.e.* the physical density of the protein-coated NP), and, consequently, the nominal and effective doses are given by Eq.s (S5) and (S6), respectively:

$$M_{nom}(x) = M(x,0) = \int M(d_p; x, 0)\partial d_p \tag{S5}$$

$$M_{eff}(t) = M(0,t) = \int M(d_p; 0, t)\partial d_p \tag{S6}$$

- $D_{diff}$ (m$^2$ s$^{-1}$) is the isotropic diffusion coefficient of each NP with diameter $d_p$, defined according to Eq. (S6) (Stokes-Einstein's equation):

$$D_{diff}(d_p) = \frac{RT}{3N_A\pi\mu d_p} \tag{S7}$$

with $R$ (J mol$^{-1}$ K$^{-1}$) the universal gas constant, $T$ (K) the temperature of the culture medium, $N_A$ (mol$^{-1}$) the Avogadro's number and $\mu$ (Pa s) the dynamic viscosity of the culture medium;

- $v_{sed}$ (m s$^{-1}$) is the steady-state sedimentation velocity of a NP of effective diameter $d_p$, which is given by Stokes' equation:

$$v_{sed}(d_p) = \frac{g(\rho_{eff} - \rho_f){d_p}^2}{18\mu} \tag{S8}$$

where $g$ (m s$^{-2}$) is gravitational acceleration and $\rho_f$ (kg m$^{-3}$) is the density of the culture medium.

The dissolution rate (Eq. (S3)) can simply be treated as a constant, holding across all NP diameters. In the case of the DG model, which accounts for dissolution considering ${R_{diss}}^*$ (s$^{-1}$), *i.e.* the fraction of the initial number surface density of NPs with a specific diameter $d_p$ dissolving per unit time, Eq. (S3) is a first order rate equation [1]:

$$R_{diss}(d_p;x) = -N(d_p;x,0){R_{diss}}^* \tag{S9}$$

On the other hand, the ISD3 implements dissolution as a surface area-driven phenomenon based on a NP-specific kinetic model. This description can be more accurate but less general than that provided in Eq. (S9). It does however require an in-depth characterization of dissolution kinetics for determining an analytical formulation of the associated rate suitable for the NP of interest [2].

Combining the rate equations introduced so far results in a single PDE, which completely describes NP dynamics in protein-enriched aqueous media:

$$\frac{\partial N(d_p;x,t)}{\partial t} = D_{diff}(d_p)\frac{\partial^2 N(d_p;x,t)}{\partial x^2} - v_{sed}(d_p)\frac{\partial N(d_p;x,t)}{\partial x} - \frac{\partial}{\partial d_p}\left(N(d_p;x,t)\frac{\partial d_p}{\partial t}\right) \tag{S10}$$

Both models in DosiGUI solve the PDE in Eq. (S10) with respect to $N$ for each existing NP size $d_p$, exploiting different numerical approaches [1], [2].

### SI3 – Analytical details about the bottom boundary condition

The adsorption of NPs on the cell monolayer which resides onto the set-up bottom is differently modelled by DG and ISD3. Further analytical specifications regarding this aspect are provided here.

#### *DG model*

The DG model accounts for a sticky bottom by implementing a Langmuir adsorption kinetics (Eq. (1) in the main text). The instantaneous adsorption rate depends on the dissociation constant $k_D$, which is the parameter to be identified and set for modulating what we called stickiness index [1]. In addition, note that the molar concentration $[NP](0,t)$ accounts for ENPs of all existing sizes $d_p$ and can be immediately derived from the dependent variable of the model (*i.e.* the number surface density of NPs, $N$) as:

$$[NP](0,t) = 10^{-3}\frac{\int N(d_p;0,t)\partial d_p}{N_A \Delta x} \qquad \text{(S11)}$$

where $\Delta x$ (m) is the spatial resolution of simulations (*i.e.* the height of the simulation compartment) and the multiplying factor $10^{-3}$ is needed for converting from $m^3$ to L.

#### *ISD3 model*

ISD3 manages the characteristics of the bottom by means of a typical mixed boundary condition for the PDE in Eq. (S9) [2]. We have purposely modified the equation describing this boundary condition (Eq. (2) in the main text), by parametrizing it with respect to $K$ (m), which determines the stickiness index of the cell monolayer (Eq. (S12)).

$$N\left(d_p;0,t\right) + K\frac{\partial N(d_p;0,t)}{\partial x} = 0 \qquad \text{(S12)}$$

Thus, if $K \rightarrow 0$, Eq. (S11) expresses a typical Dirichlet boundary condition ($N(d_p; 0, t) = 0$), corresponding to a totally instantaneous adsorptive behaviour. On the other hand, if $K \rightarrow \infty$, a Neumann boundary condition results ($\frac{\partial N(d_p;0,t)}{\partial x} = 0$), implying a purely reflective bottom boundary (*i.e.* no flux condition).

**SI4 – Statistical details about correlation analyses performed**

Correlation analysis was carried out for both model validation and identification of parameters modulating the cell layer stickiness. In particular, the regression line which best fits the trend of $\varphi_m^{exp}$ (or $\varphi_a^{exp}$) as a function of $\varphi_m^{sim}$ (or $\varphi_a^{sim}$) was determined with a confidence interval of 95% for each ENP and nominal dose. For improving its robustness, the fitting procedure was weighted according to the inverse of the squared standard deviation of each experimental measurement. DosiGUI predictions were considered as validated for the specific ENP (or the value of $k_D$ or $K$ giving the predictions considered was assumed as a suitable candidate for the specific ENP, cell type and nominal dose) on the basis of the following criteria:

- Pearson's coefficient $(r) \geq 0.8$ (*i.e.* measurements and predictions are significantly correlated);
- $p \geq 0.05$ for the extra sum-of-squares F-test on the slope of the regression line, having the null hypothesis that the best-fit value of the parameter is not statistically distinguishable from 1 (*i.e.* numerical values of predictions do not significantly differ from measurements).

Note that, since these are necessary but not sufficient conditions singularly, both criteria have to be satisfied, starting for each nominal dose considered.

For parameter identification, the most suitable value of $K$ or $k_D$ was determined as that maximizing both the computed statistics (*i.e.* the Pearson's coefficient $r$ and the p-value $p$ of the extra sum-of-squares F-test). If slightly different values were identified for the same ENP starting from different nominal doses, the overall optimal value of $K$ or $k_D$ ($K_{opt}$ or $k_{D,opt}$) for the ENP was defined as:

$$K_{opt} = \sum_i w_i K_i \tag{S13}$$

where $i$ indices nominal doses, $K_i$ (m) is the stickiness which optimizes correlation statistics for the i-th nominal dose, and $w_i = r_i p_i / \sum_i r_i p_i$ denotes the weighting for $K_i$ depending on $r$ and $p$ of the associated regression line. For instance, Eq. (S13) reports the case of the ISD3 model; the same rationale held for the definition of $k_{D,opt}$ in the DG model.

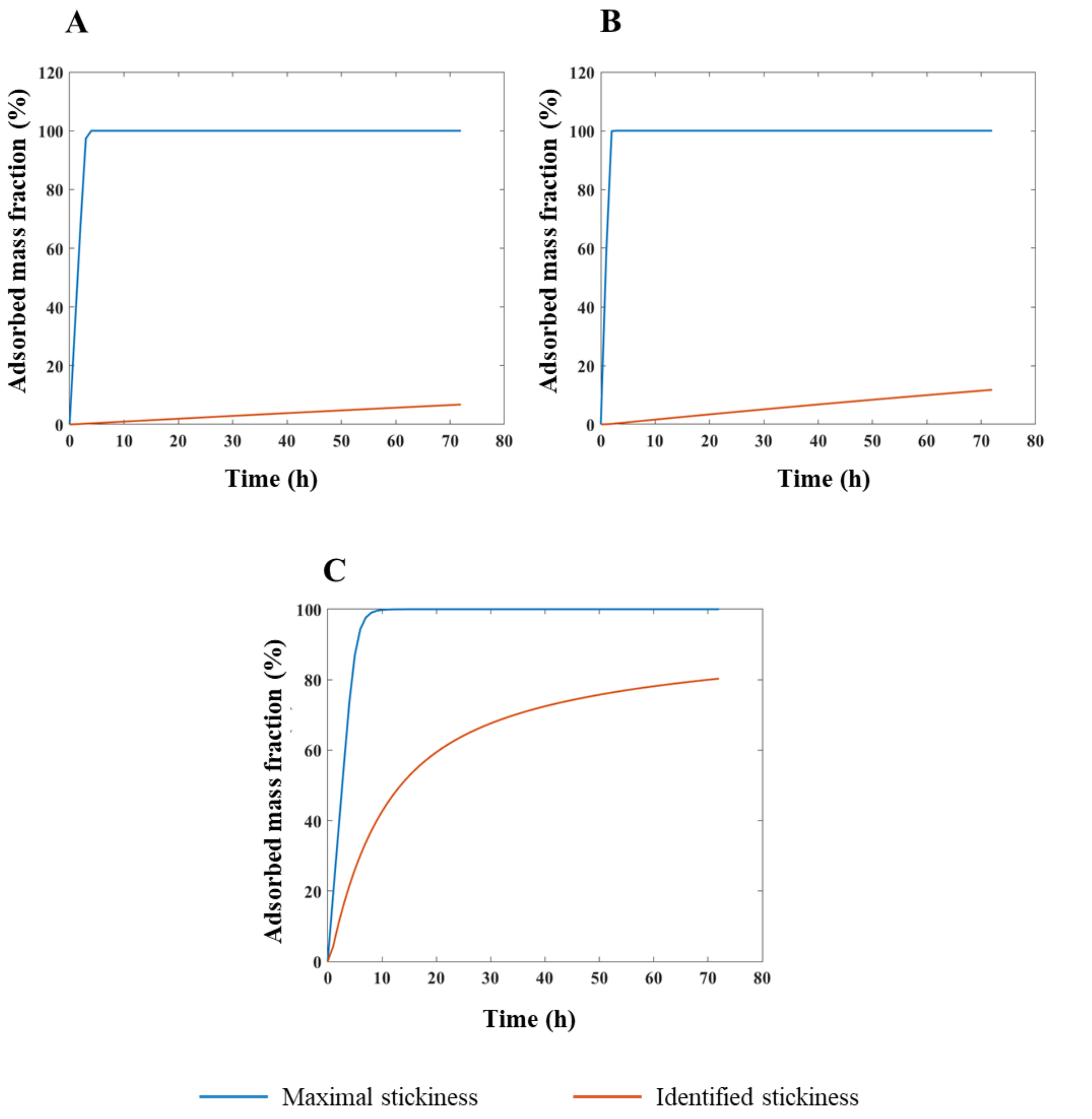


Figure S1. Predictions of mass fraction over time adsorbed by HepG2 cells expressed as a percentage of the total amount of ENP mass in the suspension, obtained running ISD3 within DosiGUI and assuming a nominally administered dose of 78.1 μg/cm$^2$. Comparison between the case of a maximally sticky bottom (blue line) and the identified stickiness index of HepG2 monolayers (orange line) for: **A)** $TiO_2$ (NM-105), **B)** $CeO_2$ (NM-212) and **C)** $BaSO_4$ (NM-220). Note that, unlike the DG model, setting a maximal stickiness in the ISD3 model implies that the initially administered mass of each ENP is completely adsorbed on the bottom in few hours. This is because the DG model simulates surface occupancy and saturation whereas ISD3 does not impose such as surface condition and thus the boundary acts like a sink.

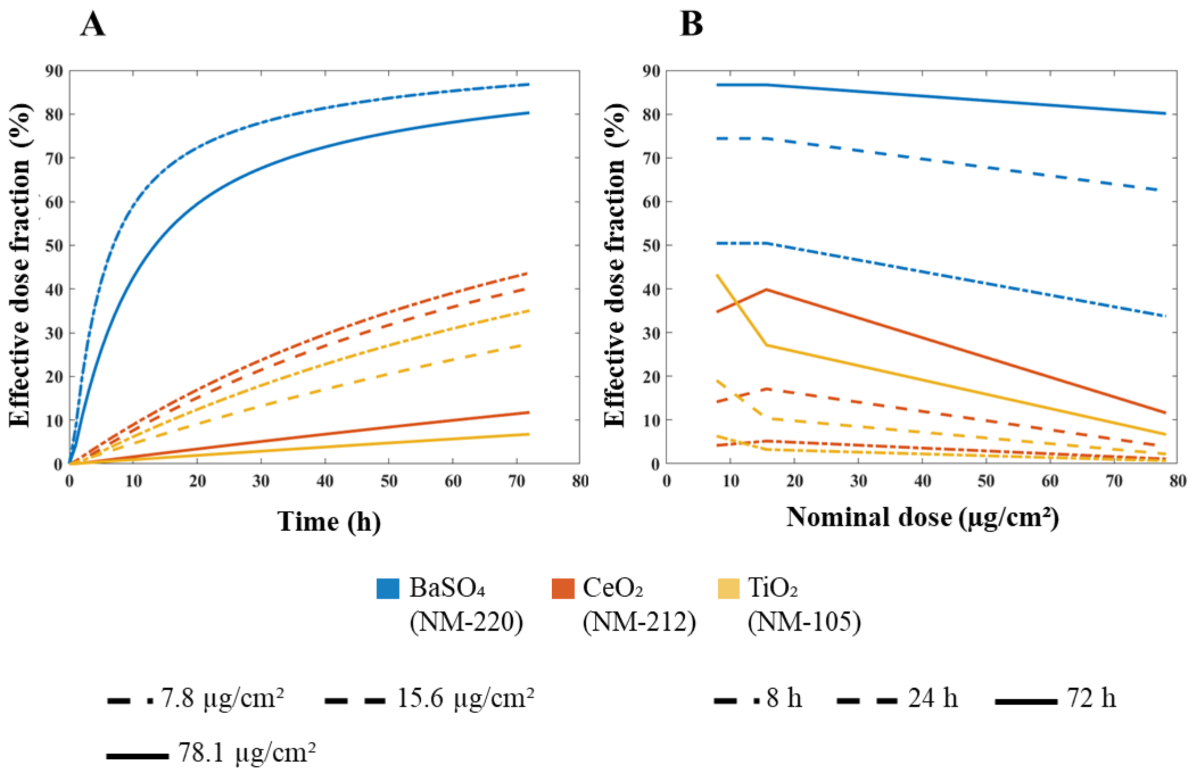


Figure S2. Predictions for the effective dose fraction (*i.e.* effective dose/nominal dose, percentage) delivered to HepG2 cells, obtained running ISD3 within DosiGUI. **A)** Effective dose fraction versus exposure time. **B)** Effective dose fraction versus nominal dose. Note that, for $BaSO_4$ (NM-220), only two curves are visible because the effective dose fraction profiles over time perfectly overlap starting from nominal doses of 7.8 and 15.6 µg cm$^{-2}$.

Table S1. Fundamental input parameters of simulations performed for model validation within DosiGUI. Physicochemical properties of ENPs are from [3].

| Parameter | Assigned value |
|---|---|
| Medium height ($h$) | 50 mm |
| Medium density ($\rho_f$) | 0.9995 g mL$^{-1}$ |
| Medium dynamic viscosity ($\mu$) | 0.81 mPa s |
| Medium temperature ($T$) | 37 °C |
| ENM nominal dose ($M_{nom}$) | 5, 25, 50 µg cm$^{-2}$ |
| Bottom boundary stickiness | $k_D$ = 0 mol L$^{-1}$, $K$ = 0 m |
| Total exposure time ($\tau$) | 24 h |
| Spatial resolution ($\Delta x$) | 0.5 mm |
| Time resolution ($\Delta t$) | 60 s |
| Dissolution constants | Dissolution disabled |
| NM-105 size distribution | Log-normal<br>Mean: 464.5 nm, Standard deviation: 2.6 nm |
| NM-212 size distribution | Log-normal<br>Mean: 243.3 nm, Standard deviation: 3.0 nm |
| NM-220 size distribution | Log-normal<br>Mean: 111.9 nm, Standard deviation: 4.5 nm |
| NM-105 density ($\rho$) | 4.23 g mL$^{-1}$ |
| NM-212 density ($\rho$) | 7.21 g mL$^{-1}$ |
| NM-220 density ($\rho$) | 4.49 g mL$^{-1}$ |
| NM-105 effective density ($\rho_{eff}$) | 1.41 g mL$^{-1}$ |
| NM-212 effective density ($\rho_{eff}$) | 2.21 g mL$^{-1}$ |
| NM-220 effective density ($\rho_{eff}$) | 2.03 g mL$^{-1}$ |

Table S2. Fundamental input parameters of simulations performed for identifying parameters determining the stickiness index in both DG and ISD3. Physicochemical properties of ENPs are from [3].

| Parameter | Assigned value |
|---|---|
| Medium height ($h$) | 3.125 mm |
| Medium density ($\rho_f$) | 0.9995 g mL$^{-1}$ |
| Medium dynamic viscosity ($\mu$) | 0.81 mPa s |
| Medium temperature ($T$) | 37 °C |
| ENM nominal dose ($M_{nom}$) | 25, 50, 250 μg cm$^{-2}$ |
| Bottom boundary stickiness | $k_D \in [1\times10^{-12}; 1\times10^{-7}]$ mol L$^{-1}$<br>$K \in [1\times10^{3}; 1\times10^{10}]$ m |
| Total exposure time ($\tau$) | 72 h |
| Spatial resolution ($\Delta x$) | 0.05 mm |
| Time resolution ($\Delta t$) | 1 s |
| Dissolution constants | Dissolution disabled |
| NM-105 size distribution | Log-normal<br>Mean: 464.5 nm, Standard deviation: 2.6 nm |
| NM-212 size distribution | Log-normal<br>Mean: 243.3 nm, Standard deviation: 3.0 nm |
| NM-220 size distribution | Log-normal<br>Mean: 111.9 nm, Standard deviation: 4.5 nm |
| NM-105 density ($\rho$) | 4.23 g mL$^{-1}$ |
| NM-212 density ($\rho$) | 7.21 g mL$^{-1}$ |
| NM-220 density ($\rho$) | 4.49 g mL$^{-1}$ |
| NM-105 effective density ($\rho_{eff}$) | 1.41 g mL$^{-1}$ |
| NM-212 effective density ($\rho_{eff}$) | 2.21 g mL$^{-1}$ |
| NM-220 effective density ($\rho_{eff}$) | 2.03 g mL$^{-1}$ |
